# Transmembrane domains of fusion proteins promote stalk formation by inducing membrane disorder

**DOI:** 10.1101/2025.08.14.670253

**Authors:** Katharina C. Scherer, Chetan S. Poojari, Jochen S. Hub

## Abstract

Membrane fusion is a fundamental process involved in exocytosis, fertilization, or cell entry by enveloped viruses. Membrane fusion is facilitated by fusion proteins, which are anchored in membranes by helical transmembrane domains (TMDs). Previous studies showed that TMD variations may alter the fusion efficiency, suggesting that TMDs are not merely passive anchors, however the mechanism by which TMDs drive fusion is not well understood. We used high-throughput coarse-grained molecular dynamics simulations and free energy calculations to quantify effects of TMDs on the formation of the first fusion intermediate, that is, of a fusion stalk. We analyzed five physiologically relevant TMDs derived from viral fusion proteins and the SNARE complex embedded in various lipid environments. We find that the addition of TMDs favors stalk formation by typically 10 to 30 kJ/mol in a concentration-dependent manner. Using helices with sequences R_2_L_n_R_2_ (*n* = 6*, . . .,* 26), we find that negative hydrophobic mismatch between the TMD and the membrane core strongly promotes fusion. Analysis of the lipid tail order parameters of annular lipids revealed a strong correlation between stalk stabilization and induced lipid disorder. Together, our findings suggest that TMDs actively contribute to membrane fusogenicity by locally perturbing the membrane order.

## Introduction

Membrane fusion is a fundamental process in cell biology, playing a crucial role in events such as the entry of enveloped viruses during infection, exocytosis, intracellular cargo trafficking, and fertilization.^1,2^ Despite these diverse contexts, membrane fusion proceeds via a common pathway, involving intermediate non-bilayer conformations. The fusion process starts with two membranes in close proximity. Overcoming hydration repulsion forces enables the formation of a fusion stalk structure with a hydrophobic connection between the proximal leaflets.^3^ Expansion of the hourglass-shaped stalk structure leads to the hemifusion diaphragm, at which point a fusion pore may form. These intermediate conformations are separated by energy barriers that need to be overcome for a successful fusion event.^4^ Here, stalk formation and fusion pore opening have been proposed to constitute the two main energy barriers.^5^ A complex fusion machinery, consisting of specific fusion proteins, drives fusion by helping to overcome or by lowering the free energy barriers.^6,7^

Membrane fusion is facilitated by fusion proteins. The SNARE protein family is involved in most types of intercellular membrane fusion. ^8^ For the infection of enveloped viruses, species-specific viral fusion proteins located on the viral surface are the main drivers of viral entry via fusion with host membranes. ^7^ Fusion proteins of the SNARE machinery as well as viral fusion proteins are anchored in the membrane by a single helical protein stretch, the so-called transmembrane domain (TMD).^9^ Several lines of evidence have shown that TMDs serve not merely as membrane anchors but play an active role during fusion (reviewed in Refs. 9–11).

Experimental studies showed that replacing the SNARE TMD with fatty acid tails results in poor fusion efficiency.^12–14^ In contrast, isolated TMD mimics of the vesicular stomatitis virus G-protein^15,16^, as well as SNARE TMD mimics^17^ can enhance fusion between liposomes. Mutations introducing *β*-branched helix-destabilizing amino acids into TMDs have been found to promote fusion in SNARE-mediated as well as viral fusion. ^15,17–20^ Indeed, SNARE TMDs and viral fusion proteins are enriched in *β*-branched amino acids such as isoleucine, valine, and glycine. ^17,18,21^ This enrichment increases backbone flexibility, facilitates transient unfolding in solution, and introduces kinks into the helical structure.^22,23^ Such flexibility of TMDs is believed to be crucial in promoting fusion, since increased flexibility in TMD mimics has been proposed to aid bilayer dehydration,^22^ and has been found to enhance lipid tail splay,^24^ a process considered the initiation of stalk formation. ^25–27^

Additionally, computational studies simulating membrane fusion have investigated the role of TMDs. In simulations of SNARE-mediated vesicle fusion, Risselada *et al.* observed that the fusion stalk consistently forms near SNARE TMDs, likely due to TMD-induced lipid packing disruption that facilitates initial lipid bridges between the fusing vesicles.^28^ They further concluded that this effect arises from intrinsic TMD properties rather than from mechanical stress transmission along the SNARE complex. Similarly, intrinsic features of model SNARE TMDs, such as their length, were found to influence the time required to initiate fusion between vesicles in coarse-grained simulations. ^29^ Furthermore, Smirnova *et al.* found that SNARE TMDs in fusing membranes reduce the free energy cost of stalk formation, and they speculated that this effect arises from the TMD length being well-suited to the thinned membrane region around the stalk structure.^30^ These findings suggest that TMDs play a key role in overcoming the first major energy barrier in fusion. However, a comprehensive mechanistic understanding of how TMDs from different fusion proteins promote stalk formation remains elusive.

We used molecular dynamics (MD) simulations and free energy calculations to quantify the effects of TMDs on stalk formation. Across five TMDs from viral fusion proteins and the SNARE complex, we found that TMDs reduce the free energy cost of stalk formation in a concentration-dependent manner. Using simplified model TMDs with the sequence R_2_L*_n_*R_2_ (*n* = 6, · · ·, 26) embedded in membranes composed of different lipids, we show that the stalk-stabilizing effect correlates with the hydrophobic mismatch between TMD and membrane core. Furthermore, across all simulated TMDs and lipid compositions, we find that the stalk-stabilizing effect of TMDs correlates with the TMD-induced reduction in the lipid tail order parameter. Notably, kinked TMDs favor stalk formation more strongly than straight helices. Together, our results suggest that TMDs generally favor stalk formation by inducing disorder in lipid tail packing.

## Methods

### Simulation setup and parameters

Simulation setup resembles previous work by Poojari *et al.*.^31^ Unbiased MD simulations were carried out with Gromacs^32^, versions 2020.5 and 2019.6. Simulations with harmonic restraints along the reaction coordinate *ξ*_ch_ were carried out with a modification of Gromacs 2018.8, ^31^ available on GitLab (gitlab.com/cbjh/gromacs-chain-coordinate). The interactions were described with the coarse-grained Martini^33^ force field, version 3.0.beta.3.2, except the data from bilayers with multiple TMDs shown in Fig. 1C/D, for which Martini version 3.0.0 was used.

**Figure 1:**
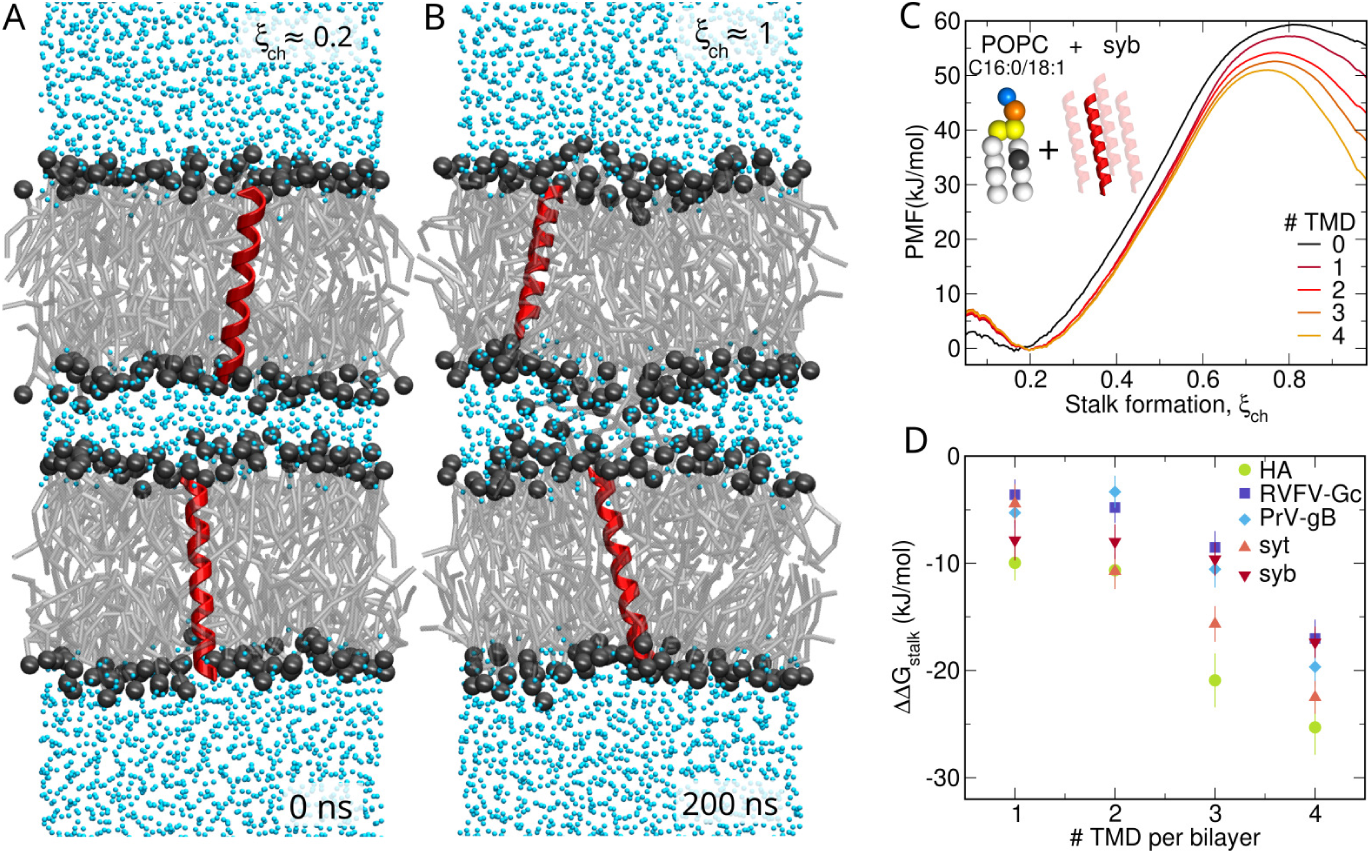
(A/B) Martini simulation system of POPC bilayers with one inserted TMD from the fusion protein Gc of Rift Valley fever virus per bilayer and 12 water beads per nm^2^ in the proximal water compartment. Representative frames of (A) flat separated membranes and (B) the stalk structure. (C) PMFs of stalk formation for POPC bilayers with zero to four TMDs from synaptobrevin (syb) per bilayer. (D) Change in stalk free energy ΔΔG_stalk_ relative to pure lipid bilayers versus number of TMDs per bilayer for TMDs from influenza virus hemagglutinin (HA), Rift Valley fever virus Gc (RVFV-Gc), pseudorabies virus glycoprotein B (PrV-gB) as well as for syntaxin (syt) and synaptobrevin (syb) from the SNARE complex.

The simulation system was composed of two lipid bilayers stacked on top of each other separated by two water compartments, one between the membranes (proximal water) and one surrounding the double membrane system across the periodic boundary in *z*-direction (distal water, Fig. 1A/B). To set up the double-membrane system, first a single lipid bilayer with 64 lipids per leaflet was built with Insane. ^34^ For systems with multiple TMDs (Fig. 1C/D), the bilayer was built within a fixed area of 100 nm^2^. Membranes were built with one of the following lipids, according to the Martini nomenclature: DBPC, DGPC, DNPC, DOPC, DPPC, DXPC, PAPC, PEPC, PGPC, PIPC or POPC (Supplementary Tab. S1). The bilayer was hydrated and a first energy minimization and equilibration for 20 ns was performed. Two copies of the bilayer were stacked on top of each other, whereas one bilayer was flipped by 180*^◦^*. The degree of hydration between the proximal leaflets was set by the number of water beads per area of the simulation box in *x*-*y*-plane, thereby controlling the head group distance of the proximal monolayers in *z*-direction. We used 8 water beads per nm^2^, except for the data in Fig. 1C/D, for which 12 water beads per nm^2^ were used. The simulation box was enlarged in *z*-direction to fully hydrate the distal leaflets with 15 water beads per lipid. The double membrane system was equilibrated for 20 ns.

Structures of TMDs were generated with PyMol.^35^ The martinize script^36^ was used to convert the TMD to the Martini representation. The amino acid sequences of the TMDs used in this work are listed in Supplementary Table S2. To induce a kink in the LV16, LV20, or L16 TMDs, the dihedral angle between four backbone beads at the center of the helix were modulated between −120*^◦^* and 120*^◦^*.

Flat-bottomed position restraints (Fb-posres) were applied to water beads to prevent water permeation across the membranes and, thereby, maintain a constant degree of hydration between the proximal leaflets. To this end, the center of the simulation box in *z*-direction was used as reference position. Attractive Fb-posres with a thickness of the flat region of *z*_fb_ = (*z*_u_ − *z*_l_)*/*2 − 0.5 nm were applied to water in the proximal compartment, where *z*_u_ and *z*_l_ is the center of mass of the upper and lower bilayer. Repulsive Fb-posres with a thickness of the flat region *z*_fb_ = (*z*_u_ − *z*_l_)*/*2 + 0.5 nm were applied to water in the distal compartment. For bilayer systems with TMDs, Fb-posres were applied to the central backbone atom of the helices to keep the TMD in a transmembrane conformation and to avoid that helices would occasionally flip to the membrane surface. ^37^ Here, a thickness of 0.2 nm was used. These Fb-posres allowed unrestricted movements of the TMDs in the *x*-*y*-plane, but movements in *z*-direction were restricted to the flat-bottomed region. The force constant of all Fb-posres was set to 500 kJ mol*^−^*^1^ nm*^−^*^2^. The double-membrane system was kept centered at the simulation box by applying center of mass pulling in *z*-direction on all lipids with the box center as reference point. Thereby, any drift of the double-membrane system was prevented to ensure that the reference coordinates of the Fb-posres on the TMDs remain at the membrane centers.

The integration time step was set to 18 fs. The Verlet-neighborlist scheme was applied. The cutoffs for the Lennard-Jones and Coulomb potential were set to 1.1 nm. Simulations were carried out in the NpT ensemble. The reference temperature was set to 310 K and controlled through velocity rescaling (*τ* = 0.5 ps) with separate temperature coupling groups for the lower and upper bilayers, the lower and upper TMDs, and for the proximal and distal water compartments. Berendsen pressure coupling was semi-isotropically applied with a reference pressure of 1 bar, a compressibility of 3×10*^−^*^5^ bar*^−^*^1^, and a time constant of 6 ps.

### Reaction coordinate and free energy calculations

To drive the transition between the flat membranes and the stalk structure, the chain reaction coordinate *ξ*_ch_ originally applied for water pore formation in lipid membranes was adapted for the stalk formation process.^31,38^ The coordinate *ξ*_ch_ is defined through a cylinder with radius *R*_cyl_ = 1.2 nm divided into *N_s_* slices of the thickness *d* = 1 Å. The cylinder is placed at the center (in *z*-direction) between the two flat bilayers spanning the proximal water compartment completely and the headgroup regions of the proximal leaflets. *ξ*_ch_ is given by the fraction of slices that are occupied by lipid tail beads:

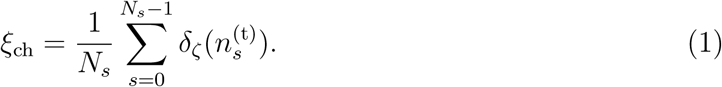

Here, 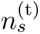 denotes the number of tail beads in slice *s* and *δ_ζ_* is a continuous approximation to an indicator function and takes *δ_ζ_* = 0 for empty slices (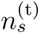 = 0) and *δ_ζ_* ≈ 1 for filled slices (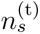 ≥ 1). The parameter *ζ* defines the degree to which a slice is filled upon the addition of the first apolar bead, here set to *ζ* = 0.75. *ξ*_ch_ quantifies the connectivity between two compartments, here between the hydrophobic cores of two opposing bilayers. By pulling along *ξ*_ch_, the cylinder is filled slice by slice with lipid tail beads, leading to a hydrophobic connection between the two membranes and, thus, to a stalk structure. More details on *ξ*_ch_ are discussed in Ref. 31.

The number of slices *N_s_* was set, depending on the degree of hydration between the bilayers, such that *ξ*_ch_ ≈ 0.2 for flat separated membranes, implying that 20 % of the cylinder slices were filled by apolar beads. The lateral position of the cylinder is defined dynamically such that the cylinder follows the stalk as the stalk moves in the *x*-*y*-plane.

Umbrella sampling was used to calculate the potential of mean force (PMF) along *ξ*_ch_. The starting frames for umbrella windows were obtained from a constant-velocity pulling simulation, where the system was pulled along *ξ*_ch_ over 200 ns with a force constant of 3000 kJ mol*^−^*^1^. We used 19 umbrella windows, and each window was simulated for 200 ns, where the first 50 ns were omitted for equilibration. For systems with multiple TMDs, we used 24 windows and a simulation time of 1500 ns per window, where the first 1000 ns were omitted for equilibration. PMFs were computed with the weighted histogram analysis method (WHAM) using the gmx wham module of Gromacs.^39^ Errors were estimated using 50 rounds of Bayesian bootstrapping of complete histograms.

The free energy for stalk formation ΔG_stalk_ was taken from the PMF by averaging the values for *ξ*_ch_ *>* 0.96. The free energy difference ΔΔG_stalk_ due to the addition of TMDs was defined as the ΔG_stalk_ value with TMD relative to the ΔG_stalk_ value of the pure lipid bilayers.

### Additional simulation analysis

The hydrophobic mismatch between polyleucine (polyL) TMDs with sequence R_2_L*_n_*R_2_ with *n* = 6, 8, 10*, . . .,* 26 and the bilayer was defined as the difference between the length of the hydrophobic region of the TMD *l*_HR_ and the thickness of the hydrophobic core of the bilayer *t*_HC_. The hydrophobic length *l*_HR_ of the polyL TMD was defined as the distance between first and last leucine backbone bead and measured using VMD. ^40^ The thickness of the hydrophobic core *t*_HC_ was taken from density profiles of the lipid tail beads computed from pure lipid bilayer simulations and taken at the point of full width at half maximum. The order parameter of the lipid tails ⟨S_N_⟩ was calculated using the script do-order-gmx5.py provided by the Martini website. To compute ⟨S_N_⟩ for the annular lipids near the TMDs, only lipids with at least one interaction bead within a distance of 2 nm to the TMD were considered.

## Results

### TMDs from viral fusion proteins or from the SNARE complex decrease the free energy for stalk formation in a concentration-dependent manner

We simulated the transition between two flat separated lipid bilayers and the stalk structure by pulling along the chain reaction coordinate *ξ*_ch_, which quantifies the degree of connectivity between the two hydrophobic membrane cores. Here, *ξ*_ch_ ≈ 0.2 corresponds to the flat separated bilayers (Fig. 1A) and *ξ*_ch_ ≈ 1 corresponds to the stalk structure (Fig. 1B). Using umbrella sampling along *ξ*_ch_, we computed the potential of mean force (PMF, also referred to as free energy profile) to obtain the free energy cost ΔG_stalk_ for stalk formation (Fig. 1C). To quantify the effect of transmembrane domains (TMDs) on stalk formation, we computed the PMFs either for pure lipid bilayers or for bilayers containing an increasing number of one to four TMDs per bilayer. Figure 1C shows PMFs for POPC bilayers with zero to four TMDs per bilayer from synaptobrevin (syb) of the SNARE complex. The PMFs reveal that the addition of TMDs decreases both the free energy barrier for stalk nucleation (*ξ*_ch_ ≈ 0.75) as well as the free energy cost of stalk formation ΔG_stalk_ (*ξ*_ch_ ≈ 1) in a concentration-dependent manner. The ΔG_stalk_ value decreases from 55 kJ/mol for a pure POPC bilayer to 50 kJ/mol with one TMD per bilayer and down to 30 kJ/mol with four TMDs per bilayer. Hence, the addition of synaptobrevin TMDs greatly facilitates stalk formation.

To test whether TMDs from other fusion proteins influence stalk formation as well, we probed the effect of four additional physiologically relevant TMDs: three TMDs from the viral fusion proteins influenza hemagglutinin (HA), Rift Valley fever virus Gc (RVFV-Gc), and pseudorabies virus glycoprotein B (PrV-gB) as well as from syntaxin (syt) from the SNARE complex. For each TMD type, PMFs of stalk formation were computed using one to four TMDs per bilayer (Supplementary Fig. S1). We calculated the change in stalk free energy ΔΔG_stalk_ relative to the pure lipid membrane, defined as the difference in stalk free energy between systems with and without TMDs. As shown in Figure 1D, ΔΔG_stalk_ decreases with the number of TMDs across all five fusion proteins. The addition of one TMD per bilayer decreases the stalk free energy ΔG_stalk_ by 3 − 10 kJ/mol, while four TMDs decrease ΔG_stalk_ by 17 − 25 kJ/mol. Hence, TMDs from the SNARE complex as well as from viral fusion proteins stabilize the fusion stalk in a concentration-dependent manner.

The decrease in the stalk free energy upon insertion of isolated TMDs is in line with Smirnova *et al.*^30^, who used the string method and the lipid tail density as order parameter to obtain the minimum free energy pathway for stalk formation. Using the identical simulation setup, kindly provided by the authors of Ref. 30, we calculated the PMF of stalk formation along *ξ*_ch_. In qualitative agreement with Ref. 30, our PMFs demonstrate that the addition of TMDs lowers the stalk nucleation barrier and stabilizes the stalk structure (Supplementary Fig. S2). However, the ΔG_stalk_ values, obtained using umbrella sampling along *ξ*_ch_ are smaller than the values reported in Ref. 30.

### Stalk stabilization by TMDs strongly depends on hydrophobic mismatch and correlates with increased lipid tail disorder

A putative mechanism by which TMDs favor stalk formation may be related to the mismatch between the hydrophobic length of the TMDs and the hydrophobic thickness of the membrane core.^41,42^ Indeed, structural membrane rearrangements for compensating such hydrophobic mismatch, such as membrane thinning, have been hypothesized as the underlying mechanism by which isolated TMDs may favor stalk formation.^30^ To test this hypothesis quantitatively, we investigated the effect of polyleucine helices (polyL) with varying hydrophobic length on ΔG_stalk_. Since two arginine residues were located at the two termini to anchor the TMD termini to the headgroup regions, the sequence of polyL was: R_2_L*_n_*R_2_ with *n* = 6, 8, 10*, . . .,* 26. Adding these polyL helices into the bilayers resulted in pronounced negative hydrophobic mismatch for short polyL up to positive hydrophobic mismatch for long polyL (Fig. 2A).

**Figure 2:**
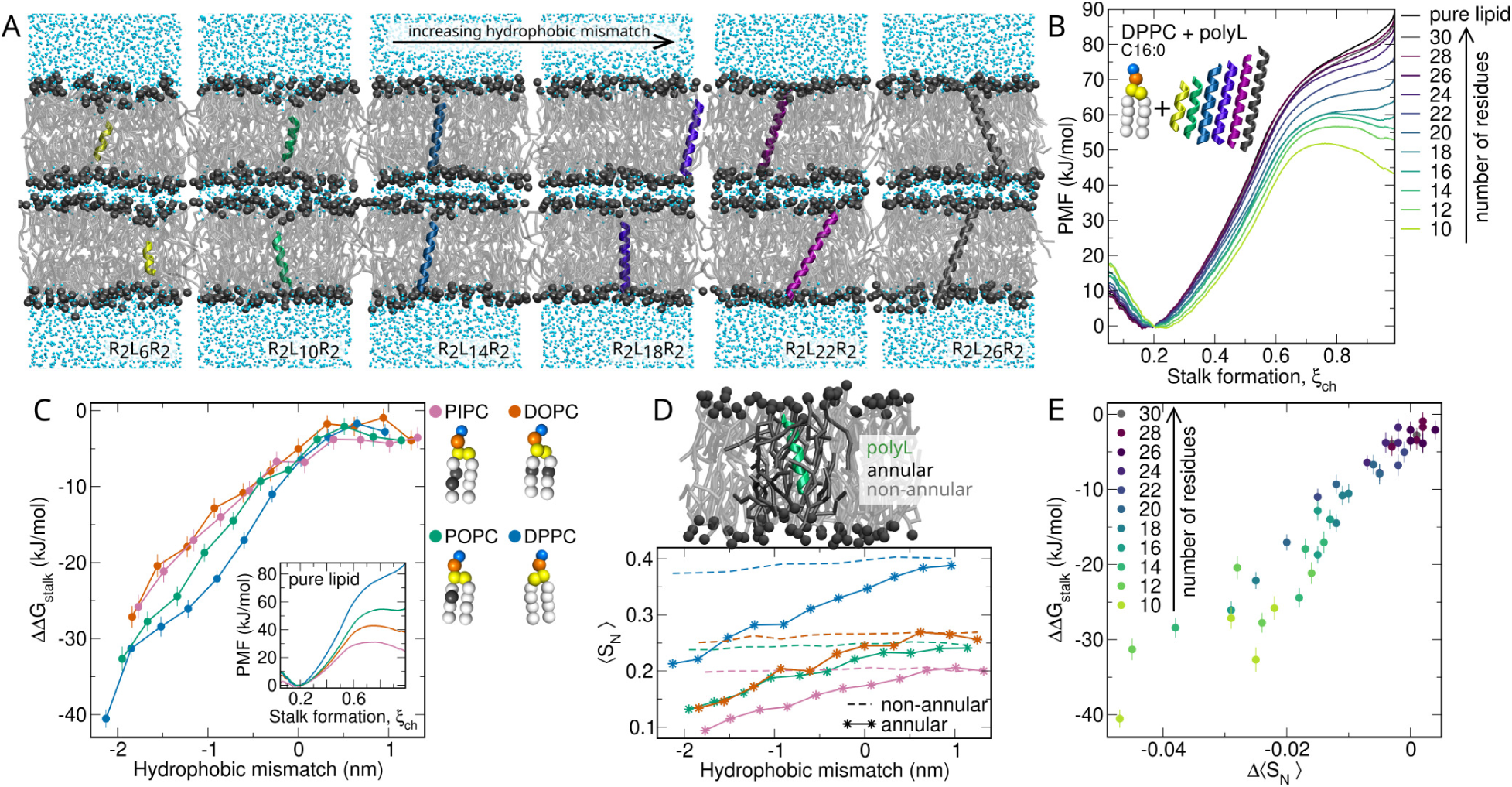
(A) Martini simulation setup of POPC bilayers with polyleucine helices (polyL) of increasing hydrophobic length and, thereby increasing hydrophobic mismatch: R_2_L*_n_*R_2_ with *n* = 6, 8, 10*, . . .,* 26. (B) PMFs of stalk formation between DPPC bilayers with polyleucine helices with increasing length (see color code). The PMF for the pure lipid bilayer is shown in black for reference. (C) Change in stalk free energy, ΔΔG_stalk_, versus hydrophobic mismatch between polyleucine helices and the membrane core for bilayers of PIPC (pink), DOPC (orange), POPC (green), or DPPC (blue). Inset: PMFs of stalk formation for pure lipid bilayers. Beads of Martini lipid models are colored as follows: hydrophobic saturated (white), hydrophobic unsaturated (grey), glycerol (yellow), phosphate (orange), choline (blue). (D) Order parameter ⟨S_N_⟩ for annular lipids (solid line) or non-annular lipids (dashed line) versus hydrophobic mismatch. (E) Change in stalk free energy ΔΔG_stalk_ versus change in order parameter Δ⟨S_N_⟩ upon insertion of polyleucine helices.

Figure 2B presents PMFs of stalk formation between two DPPC bilayers containing one polyL with increasing length per bilayer. Evidently, upon adding polyL containing ≥ 22 leucine residues, ΔG_stalk_ hardly decreases relative to the value for pure DPPC of 86 kJ/mol. In contrast, upon adding shorter polyL, ΔG_stalk_ is strongly reduced down to 43 kJ/mol for R_2_L_6_R_2_. Notably, adding polyL may change the shape of the PMF qualitatively. Stalks are unstable for pure DPPC or for systems with polyL with *n* ≥ 14 leucines, as shown by the PMF maxima at *ξ*_ch_ = 1. In contrast, in the presence of polyL with *n* ≤ 10 leucines stalks are metastable, as shown by the local free energy minima at *ξ*_ch_ = 1.

To test whether the effects of polyL TMDs depend on the type of lipid tails, we recomputed PMFs for all polyL TMDs using lipids with the same tail length as DPPC (4 beads per tail) but with increasing degree of unsaturation, namely POPC, DOPC, and PIPC (Supplementary Fig. S3). With increasing unsaturation, the lipid membranes are more disordered and the hydrophobic core is thinner compared to the membrane of saturated DPPC. Figure 2C presents the correlation between the change in stalk free energy ΔΔG_stalk_ upon insertion of polyL and the hydrophobic mismatch between the TMD and the membrane core for the four lipid types. Evidently across the four lipid types, the more negative the hydrophobic mismatch, the lower ΔΔG_stalk_, indicating increasingly favorable stalk formation. Specifically, for a hydrophobic mismatch of around −2 nm, ΔΔG_stalk_ takes values between −25 kJ/mol and −40 kJ/mol. In contrast, with positive mismatch, ΔΔG_stalk_ is larger than −5 kJ/mol in all phosphatidylcholine bilayers, demonstrating a marginal effect on stalk formation. These data demonstrate that transmembrane helices with negative hydrophobic mismatch greatly favor stalk formation.

Furthermore, the ΔΔG_stalk_ values in Figure 2C reveal that polyL takes a larger effect in the fully saturated DPPC bilayer as compared to the polyunsaturated PIPC or DOPC membranes. This indicates that the effect of transmembrane helices depends not only on the hydrophobic mismatch but also on the intrinsic order of the membranes, which is influenced by the degree of unsaturation. It has been shown before that stalk formation is facilitated between membranes of polyunsaturated lipids compared to saturated lipids (Fig. 2C Inset). ^31^ We hypothesized that the addition of a TMD into a disordered polyunsaturated membrane has a smaller effect on ΔΔG_stalk_ as compared to TMD insertion into a more ordered saturated membrane, because the TMD may influence ΔG_stalk_ by inducing disorder.

To test this hypothesis, we examined the lipid tail order. We computed the order parameter ⟨S_N_⟩ for the annular lipids, i.e. for lipids in the vicinity of the TMD. Evidently, the annular lipids exhibit increased disorder with increasingly negative hydrophobic mismatch (Fig. 2D, solid lines). Notably, the tail order of non-annular lipids in the bulk of the membrane hardly depends on the hydrophobic mismatch showing that TMDs induce lipid disorder mostly locally (Fig. 2D, dashed lines).

Additionally, we investigated the change in the order parameter averaged over all lipid tails in the bilayer upon insertion of polyL Δ⟨S_N_⟩, taken as the difference in ⟨S_N_⟩ for bilayers with or without a transmembrane helix. Remarkably, the change in lipid tail order Δ⟨S_N_⟩ strongly correlates with the change in stalk free energy ΔΔG_stalk_ (Fig. 2E). In other words, the more disorder a polyL helix induces in the lipid tails the greater the stabilization of the fusion stalk. Hence, our calculations suggest that the locally increased tail disorder is the key mechanism by which TMDs facilitate stalk formation.

### Lipid tail length and saturation determine the decrease in stalk free energy upon insertion of viral and SNARE TMDs

Since not only the TMD type but also the lipid type may influence ΔΔG_stalk_ (Fig. 2C), we systematically investigated the effect of various lipid properties on ΔΔG_stalk_. Here, we used the five TMDs as before, three TMDs from viral fusion proteins (HA, RVFV-Gc, and PrV-gB) and two TMDs from the SNARE complex (syt and syb) and added one TMD per bilayer. First, we investigated the TMD effect in phosphatidylcholine bilayers with one, two or four unsaturated lipid tail beads while keeping approximately the same tail length (Supplementary Fig. S4). As shown in Figure 3A, the TMD effect on the stalk free energy decreases with increased lipid tail unsaturation. While the change in ΔG_stalk_ is between −8 kJ/mol and −12 kJ/mol in the case of monounsaturated POPC, ΔΔG_stalk_ is between 0 kJ/mol and −6 kJ/mol for polyunsaturated PAPC lipid bilayers. As observed for polyL helices discussed above (Fig. 2C), the effect of physiological TMDs is more pronounced in bilayers with a low level of unsaturation.

**Figure 3:**
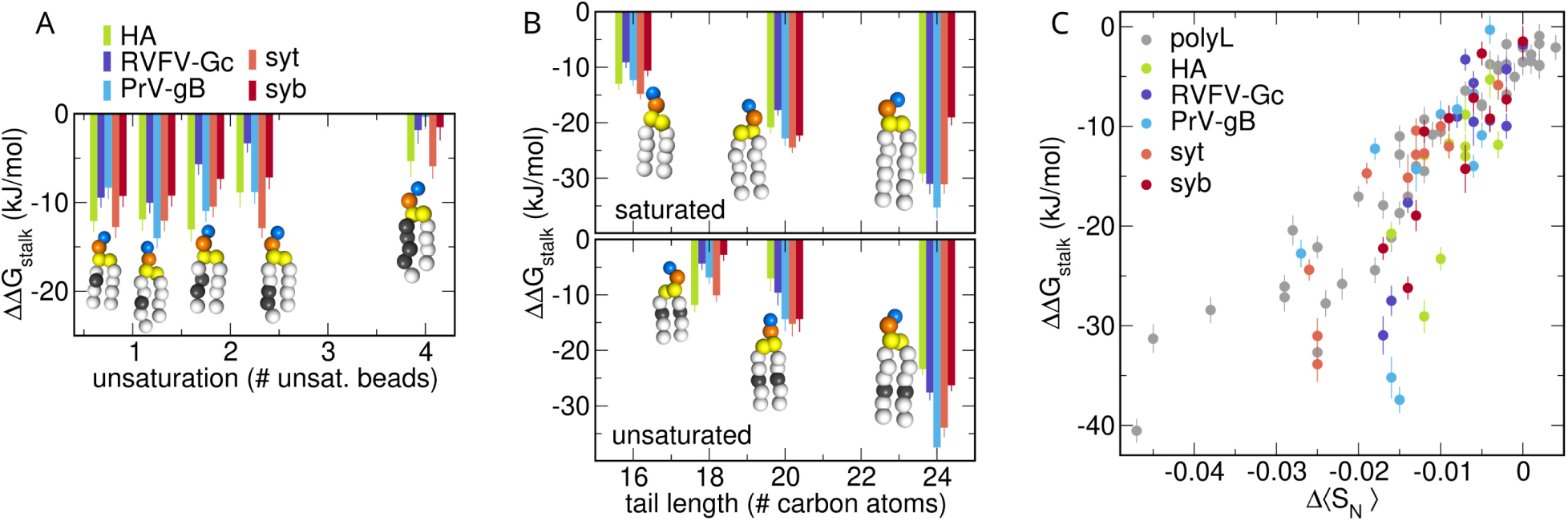
(A/B) Change in stalk free energy ΔΔG_stalk_ upon insertion of one TMD from different fusion proteins (see color code) in various lipid compositions with (A) increasing unsaturation: POPC, PGPC, PIPC, PEPC or PAPC and (B) increasing tail length: DPPC, DBPC or DXPC (top); DOPC, DGPC or DNPC (bottom). Martini beads colored as in Fig. 2. (C) Change in stalk free energy ΔΔG_stalk_ versus change in order parameter Δ⟨S_N_⟩ upon insertion of one TMD per bilayer.

Second, focusing on the tail length of lipids, Figure 3B presents ΔΔG_stalk_ upon insertion of one TMD in bilayers with four, five or six Martini beads per tail. Here, we scanned the fully saturated lipids DPPC, DBPC, or DXPC; as well as the monounsaturated lipids DOPC, DGPC, or DNPC (Supplementary Fig. S5). Regardless of the type of transmembrane peptide, the TMD effect on ΔG_stalk_ is more pronounced in membranes with longer lipid tails. Specifically, TMDs decrease ΔG_stalk_ by nearly 15 kJ/mol when inserted into bilayers with four beads per tail modeling 16 or 18 carbon atoms. The same TMDs decrease ΔG_stalk_ by nearly 37 kJ/mol when inserted into bilayers with six beads modeling 24 or 26 carbon atoms. Hence, the more negative the hydrophobic mismatch between the physiological TMDs and the lipid membranes, the more ΔG_stalk_ is reduced, in agreement with the findings on polyL helices presented above (Fig. 2).

In addition, in line with the results on polyL helices, we found that ΔΔG_stalk_ correlates with the TMD-induced disorder quantified by Δ⟨S_N_⟩ (Fig. 3C), suggesting that the physiologically relevant TMDs likewise facilitated stalk formation by inducing disorder. Few outliers from this trend, shown in Figure 3C, for which ΔΔG_stalk_ is more negative than expected from Δ⟨S_N_⟩, are taken from the exceptionally thick DXPC and DNPC membranes; hence, only for these hardly physiologically relevant membranes Δ⟨S_N_⟩ is not a precise indicator for ^ΔΔG^_stalk_.

### Kinked transmembrane helices enhance the effect on the stalk free energy

Previous experimental studies reported a correlation between fusogenicity and backbone flexibility of TMDs (reviewed in Refs. 9,10). By measuring membrane capacitance for Ca^2+^-triggered SNARE-mediated fusion, it was demonstrated that mutations in the synaptobrevin TMD to helix-stabilizing leucine reduce fusion efficiency, whereas mutations to helix-destabilizing valine or isoleucine maintain fusion efficiency relative to the wild-type TMD.^20^ Similarly, NMR spectroscopy experiments observing liposome fusion with either rigid or flexible isolated TMD mimics, where the flexibility was controlled by valine content, revealed that stalk formation and lipid tail splay is more promoted between liposomes with flexible TMDs.^24^ Furthermore, mutations in viral TMDs from glycine (GxxxG) to alanine motifs (AxxxA), decreased the rate of PEG-mediated fusion between small unilamellar vesicles.^15^ Since *β*-branched amino acids such as valine or isoleucine enhance the backbone flexibility while glycine even induces kinks in the helix^21–23^, these studies suggest that TMD flexibility is a key driver for fusion.

Since the Martini force field requires the use of secondary structure restraints, it is difficult to simulate different degrees of helix flexibility caused by *β*-branched amino acids with our setup. Instead, we tested whether bending of the TMD helix influences ΔG_stalk_. Figure 4 presents the PMFs of stalk formation between two POPC bilayers with kinked transmembrane helices LV16 with the sequence K_3_W(LV)_8_K_3_.^24^ We introduced the kink by varying one dihedral angle between four backbone beads at the center of the helix between −120*^◦^* and 120*^◦^* thereby forming angles between 85*^◦^* and 180*^◦^* between the first and the second half of the helix. The PMFs in Figure 4 revealed that bending of the helix decreases ΔG_stalk_ by nearly 25 kJ/mol as well as the stalk formation barrier at *ξ*_ch_ ≈ 0.75 by nearly 15 kJ/mol. Hence, kinked LV16 helices promote stalk formation more effectively than a straight LV16 helix. This trend is confirmed by simulations with L16 helices (sequence K_3_W(L)_16_K_3_) or LV20 (sequence K_3_W(LV)_10_K_3_) (Supplementary Fig. S6). Furthermore, the correlation between ΔΔG_stalk_ and Δ⟨S_N_⟩ obtained with these helices agrees reasonably with the correlation found for polyL (Supplementary Fig. S6), suggesting that kinked helices favor stalk formation likewise by inducing membrane disorder.

**Figure 4:**
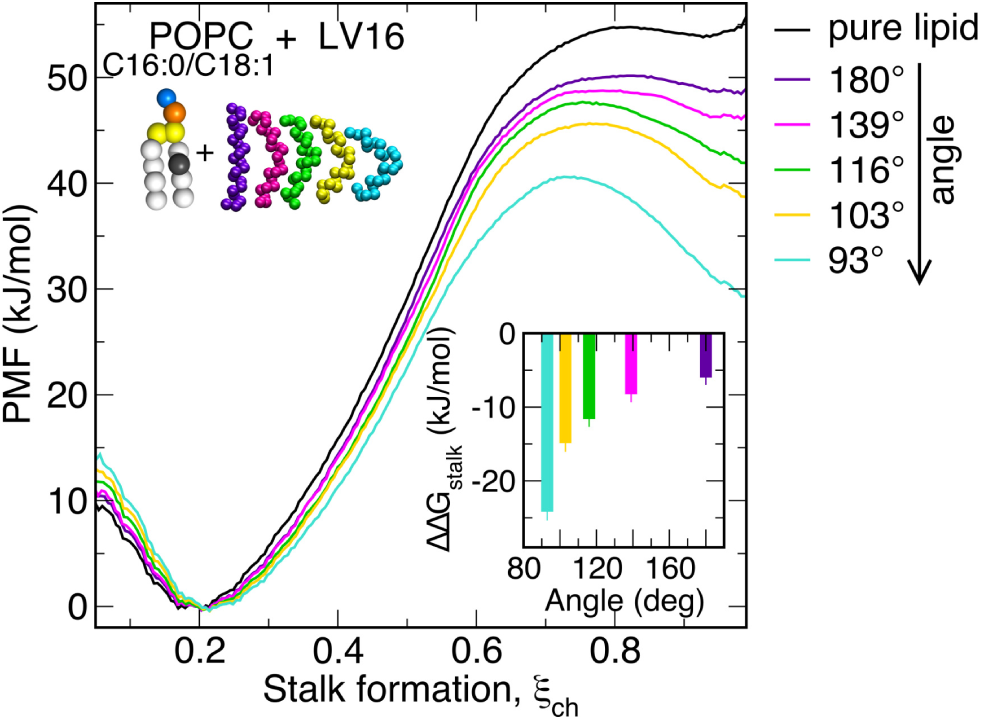
PMFs of stalk formation between two POPC bilayers with inserted LV16 (K_3_W(LV)_8_K_3_) with different bending angles. Inset: Change in stalk free energy ΔΔG_stalk_ versus bending angles.

Notably, kinked TMDs exhibit a more negative hydrophobic mismatch compared to their straight analogs. However, we observe that a straight TMD with the same hydrophobic mismatch as the most kinked LV16 TMD decreases ΔG_stalk_ to a lesser extent than the kinked helix (Supplementary Fig. S6), demonstrating that kinks contribute to the stalk formation beyond the effect of hydrophobic mismatch alone. Taken together, our simulations are supported by experiments showing that increased TMD flexibility promotes fusion. However, our simulations suggest that the underlying mechanism is not merely TMD flexibility per se, but rather the population of kinked TMD structures, which enhance lipid tail disorder.

Overall, our simulation demonstrate that TMD-induced disorder is the key factor by which TMDs favor stalk formation. We observed two mechanisms, by which TMDs enhance lipid tail disorder: either through hydrophobic mismatch or through kinked TMD structures.

## Discussion

Understanding how protein–membrane interactions control biomembrane mechanics is a vital area of research in cell biology. Amphiphilic helices or proteins containing Bin-Amphiphysin-Rvs (BAR) domains regulate membrane bending rigidity, thereby altering membrane curvature, a process crucial for cellular events such as neurotransmission and endocytosis. ^43^ Similarly, antimicrobial peptides increase membrane permeability, leading to leakage and cell death, which gives them promising potential as alternatives to conventional antibiotics.^44^ Protein crowding may affect membrane shear viscosity, thereby regulating the mobility of membrane components and influencing diffusion-limited reactions at the membrane interface.^45^ Conversely, mechanical properties of membranes such as compressibility influence transmembrane protein insertion and function, ^46^ highlighting the bidirectional role of membrane–protein interactions. In this study, we showed that TMD anchors modulate lipid ordering, promoting large-scale rearrangements into non-bilayer structures such as fusion stalks. Our data emphasizes that the TMDs of fusion proteins act as more than passive anchors; rather, they function as membrane-active peptides that modulate local membrane mechanics at their site of action.

Building on our observation that transmembrane domains introduce disorder into lipid tail packing, we hypothesize that this disruption may facilitate a local enrichment of polyunsaturated lipids. These lipids are known to favor the stalk formation.^31^ Thus, TMD-induced membrane disorder may create microenvironments that selectively recruit lipid species preferring disordered regions, thereby promoting membrane remodeling at fusion sites. This mechanism may be particularly relevant in the context of viral infection, as viruses can actively modulate host lipid synthesis, typically increasing polyunsaturated lipid content while reducing saturated lipids, potentially priming membranes for fusion events.^47,48^

The high-throughput nature of this study was possible by our efficient method for obtaining the free energy of stalk formation, based on PMF calculations along the chain coordinate (https://gitlab.com/cbjh/gromacs-chain-coordinate). This approach enabled us to compute approximately 140 PMFs to probe a wide range of physiologically relevant and model transmembrane domains across bilayers of various compositions. Only through our systematic large-scale analysis, we identified the TMD-induced lipid disorder as a unifying mechanism underlying facilitated stalk formation. The computational efficiency of the method opens the door to future investigations of more complex fusion scenarios, for instance including those involving lipid nanoparticles or ionizable lipids.

We showed that increasingly negative hydrophobic mismatch between the TMD and the hydrophobic bilayer core reduces the free energy of stalk formation, in line with previous suggestions.^30^ Notably, whereas the length of many transmembrane helices from membrane proteins often match with the membrane thickness^49^, the neuronal SNARE TMDs are shorter than the average plasma membrane thickness.^11,42^ This may be an adaptation to promote rapid and energetically efficient fusion of neuronal vesicles. Recent coarse-grained simulations of SNARE mimics suggested that fusion rates may increase with shortened TMDs. ^29^ In line with these findings, the length of TMDs from viral proteins was found to influence the entry pathway of viruses.^50^ Our results corroborate these findings by revealing that shorter TMDs with negative hydrophobic mismatch induce more disorder in lipid tails and, thereby, stabilize the stalk structure more as compared to TMDs with no or positive mismatch. Our findings furthermore align with the general observation that negative hydrophobic mismatch in transmembrane proteins promotes non-lamellar lipid phases by driving lipid rearrangements that compensate for the mismatch.^41^

We observed that kinked TMDs favor stalk formation more as compared to straight TMDs, correlated with increasingly induced lipid tail disorder. These findings rationalize the abundance of *β*-branched and helix-disrupting amino acids such as isoleucine, valine, and glycine in the TMDs from the SNARE complex and from viral fusion protein.^17,18,21,22,51^ Previous studied rationalized the effect in terms of enhanced TMD flexibility, backbone dynamics, or transient helix unfolding. However, our simulations revealed a stalk-stabilizing effect by kinked helices with fixed bending angles, i.e. in absence of intrinsic TMD flexibility. Thus, we suggest that not the TMD flexibility per se, but rather the population of kinked TMD conformers is the driver for increased lipid disorder and, consequently, for enhanced stalk formation.

The computationally efficiency of the coarse-grained Martini force field enabled the high-throughput simulations of stalk formation in this study. This leads to shortcomings since variations of head group hydration or hydrogen bonding networks owing to changing monolayer curvature are not explicitly represented.^52^ However, considering that the Martini force field often reproduces trends, we expect that trends of the stalk free energy with varying TMD length, TMD concentration or with varying degrees of lipid unsaturation are correct. Nevertheless, it will be desirable to compare the trends observed here with results from all-atom force field in future studies.

In summary, our PMF calculation showed that TMDs inserted in lipid bilayers facilitate stalk formation during membrane fusion as evidenced by a decrease in the free energy cost of stalk formation. This stabilization of the stalk was found across physiologically relevant TMDs from viral fusion proteins or from the SNARE complex in a wide range of phosphatidylcholine bilayers. By quantifying the lipid tail order parameters of the fusing membranes, we found that reduced stalk free energy correlates with the TMD-induced disorder across all systems. Thus, locally induced lipid disorder emerges as a unifying effect underlying enhanced stalk formation by TMDs. Since lipid disorder was specifically promoted by TMDs with negative hydrophobic mismatch or by kinked TMDs, such TMDs greatly favor stalk formation. This study underscores the active role of membrane anchors from fusion protein in facilitating fusion by locally altering membrane mechanics.

## Code availability statement

A modified Gromacs version that implements the chain coordinate *ξ*_ch_ along with documentation and tutorials is available at:

https://gitlab.com/cbjh/gromacs-chain-coordinate.

## Acknowledgement

This study was supported by the Deutsche Forschungsgemeinschaft (grants SFB 1027/B7 and INST 256/539-1).

