## Supporting Information PDF for "Transmembrane domains of fusion proteins promote stalk formation by inducing membrane disorder"

### Supplementary information for: Transmembrane domains of fusion proteins promote stalk formation by inducing membrane disorder

Katharina C. Scherer, Chetan S. Poojari, and Jochen S. Hub*

*Theoretical Physics and Center for Biophysics, Saarland University, Saarbrücken, Germany*

**Supplementary Table S1:** Lipid naming according Martini nomenclature, full name and atomistic equivalent.

| Martini | full name | atomistic |
|---|---|---|
| DBPC | Diarachidoylphosphatidylcholine | di–C20:0–C22:0 PC |
| DGPC | Di-gondoic-acid-phosphatidylcholine | di–C20:1–C22:1 PC |
| DNPC | Di-nervonic-acid-phosphatidylcholine | di–C24:1–C26:1 PC |
| DOPC | Dioleoylphosphatidylcholine | di–C16:1–C18:1 PC |
| DPPC | Dipalmitoylphosphatidylcholine | di–C16:0–C18:0 PC |
| DXPC | Dilignoceroylphosphatidylcholine | di–C24:0–C26:0 PC |
| PAPC | 1-stearoyl-2-arachidonoyl-phosphatidylcholine | C16:0/20:4 PC |
| PEPC | 1-stearoyl-2-eicosadienoyl-phosphatidylcholine | C16:0/20:2 PC |
| PGPC | 1-palmitoyl-2-docosenoyl-phosphatidylcholine | C16:0/20:1 PC |
| PIPC | 1-palmitoyl-2-linoleoyl-phosphatidylcholine | C16:0/18:2 PC |
| POPC | 1-palmitoyl-2-oleoyl-phosphatidylcholine | C16:0/18:1 PC |

**Supplementary Table S2:** Amino acid (aa) sequences of transmembrane domains (TMD).

| fusion protein | aa sequence of TMD |
|---|---|
| influenza hemagglutinin (HA) | WILWISFAISCFLLCVVLLGFIM |
| Rift Valley fever virus Gc (RVFV-Gc) | TILLICLYVALSIGLFFLLIYLG |
| pseudorabies Virus gB (PrV-gB) | NPFGALAIGLLVLAGLVAAFLAY |
| syntaxin (syt) | KIMIIICCVILGIIIASTIGGIFG |
| synaptobrevin (syb) | MMIILGVICAIILIIIIVYFST |
| polyleucine (polyL) | $R_2L_nR_2$, $n = 6, \ldots, 26$ |
| L16 | $K_3W(L)_{16}K_3$ |
| LV12 | $K_3W(LV)_6K_3$ |
| LV16 | $K_3W(LV)_8K_3$ |
| LV20 | $K_3W(LV)_{10}K_3$ |

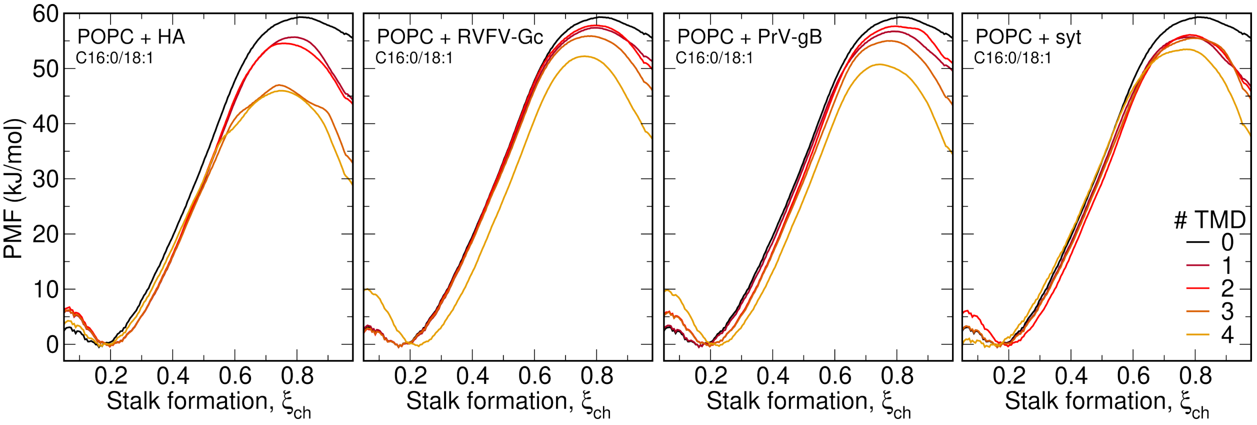


**Supplementary Figure S1:** PMFs of stalk formation for POPC bilayers with zero to four TMDs (for color code, see legend) from the following fusion proteins (from left to right): influenza virus hemagglutinin (HA), Rift Valley fever virus Gc (RVFV-Gc), pseudorabies virus glycoprotein B (PrV-gB), syntaxin (syt).

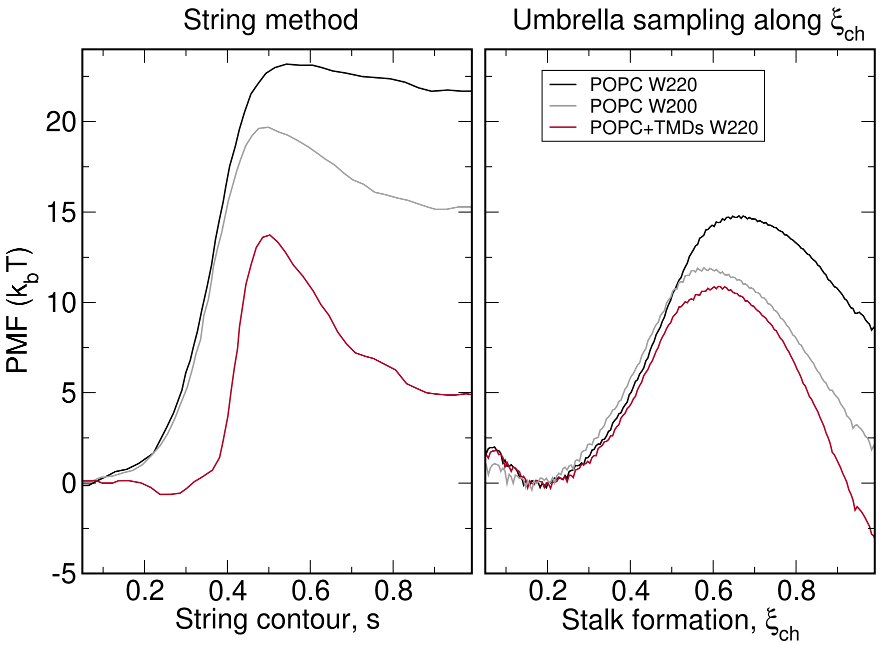


**Supplementary Figure S2:** Left: Minimum free-energy path for stalk formation between two POPC bilayers at two different degrees of hydration (W220, W200) and with inserted TMDs of the SNARE complex for the W200 system, taken from Ref. 1.
Right: PMFs computed in this work for the same simulation systems kindly provided by Smirnova *et al.*[1] obtained with umbrella sampling along the chain coordinate $\xi_{ch}$.
Effects of different degrees of hydration and the effect of the TMD agree qualitatively between the two methods. However, PMFs computed along $\xi_{ch}$ suggest smaller free energies of stalk formation as compared to previous calculations with the string method.

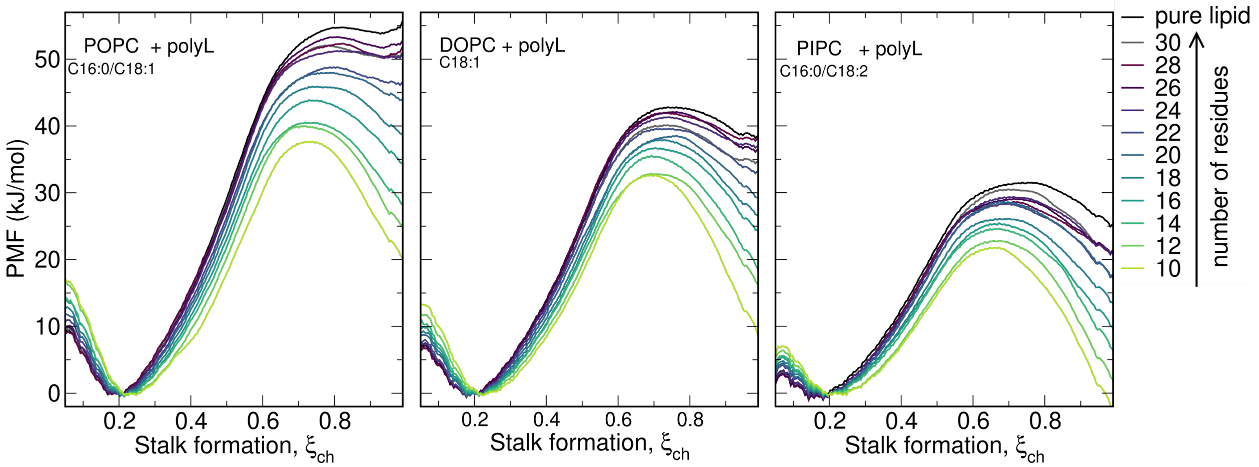


**Supplementary Figure S3:** PMFs of stalk formation between membranes of (from left to right) POPC, DOPC, or PIPC with one polyleucine helix (polyL) each with increasing hydrophobic length as controlled by the sequence $R_2L_nR_2$ ($n = 6, 8, 10, ..., 26$). The black line shows the PMF for pure lipid bilayers for reference.

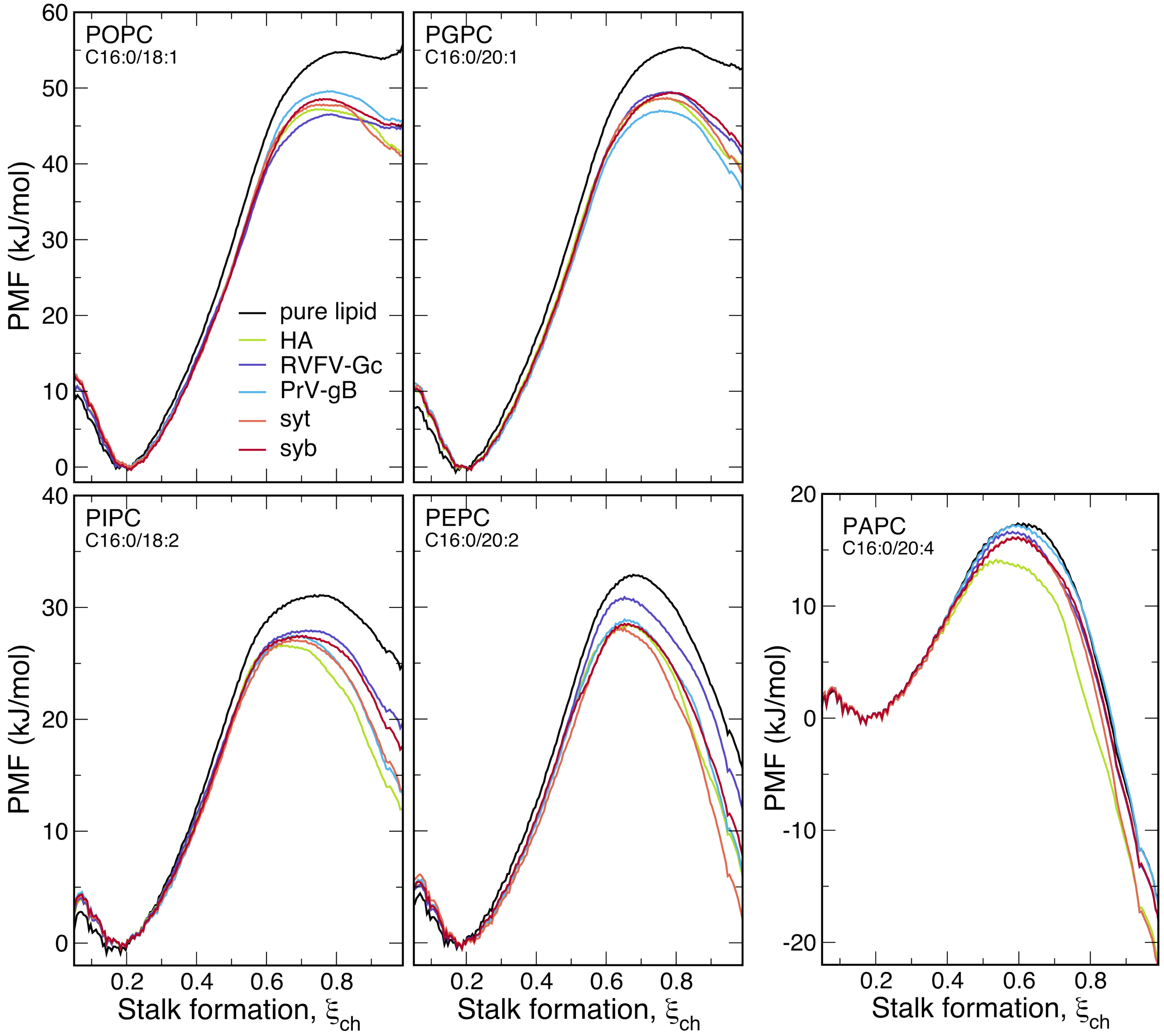


**Supplementary Figure S4:** PMFs of stalk formation between membranes composed of POPC, PGPC, PIPC, PEPC or PAPC (see labels) with one TMD from influenza virus hemagglutinin (HA, green), Rift Valley fever virus Gc (RVFV-Gc, purple), pseudorabies virus glycoprotein B (PrV-gB, blue), sytaxin (syt, orange), or synaptobrevin (syb, red). The black line shows the PMF for pure lipid bilayers for reference.

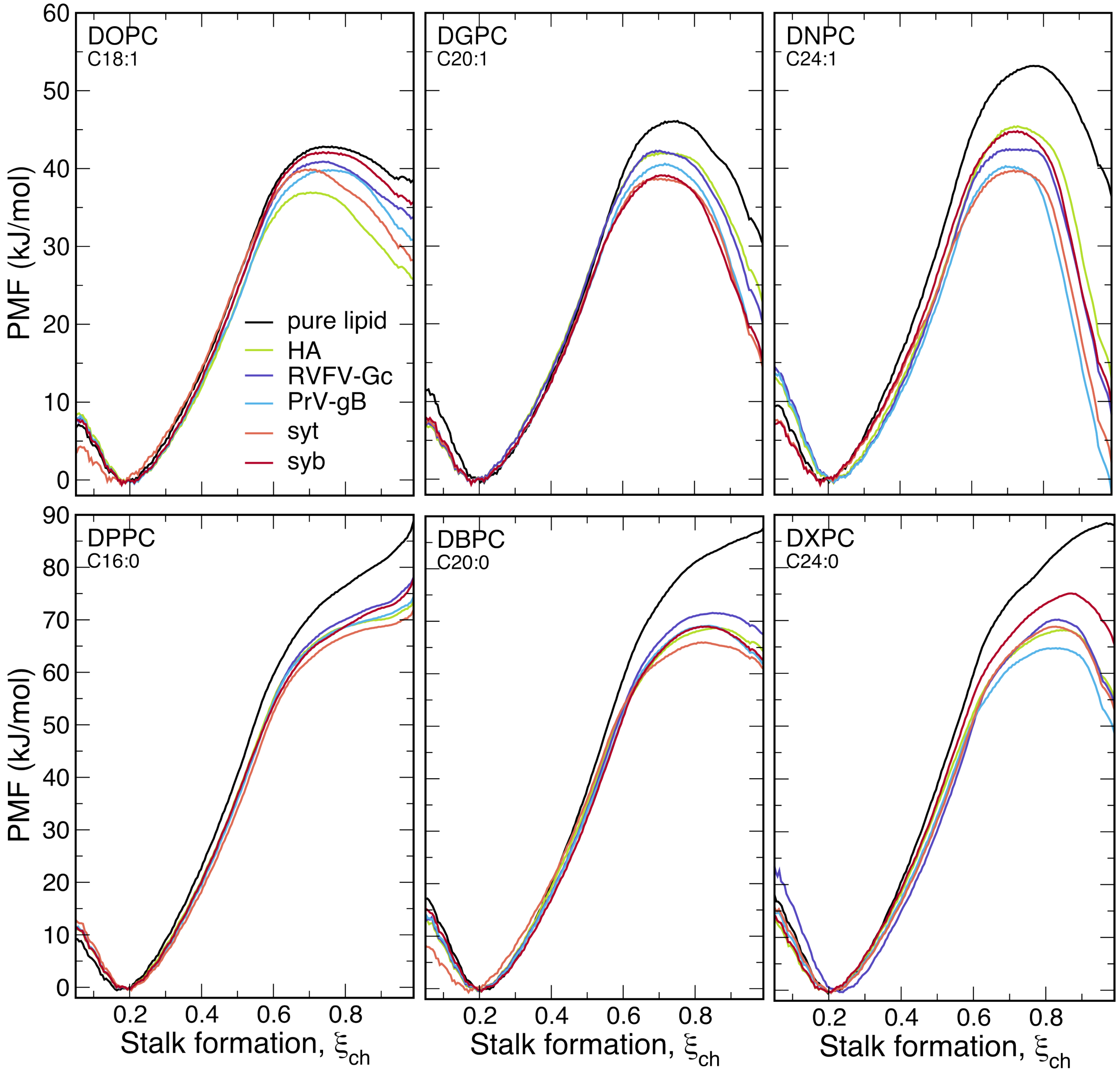


**Supplementary Figure S5:** PMFs of stalk formation between membranes composed of POPC, DOPC, DGPC, DNPC, DPPC, DBPC, and DXPC with one TMD from influenza virus hemagglutinin (HA, green), Rift Valley fever virus Gc (RVFV-Gc, purple), pseudorabies virus glycoprotein B (PrV-gB, blue), sytaxin (syt, orange), or synaptobrevin (syb, red). The black line shows the PMF for pure lipid bilayers for reference.

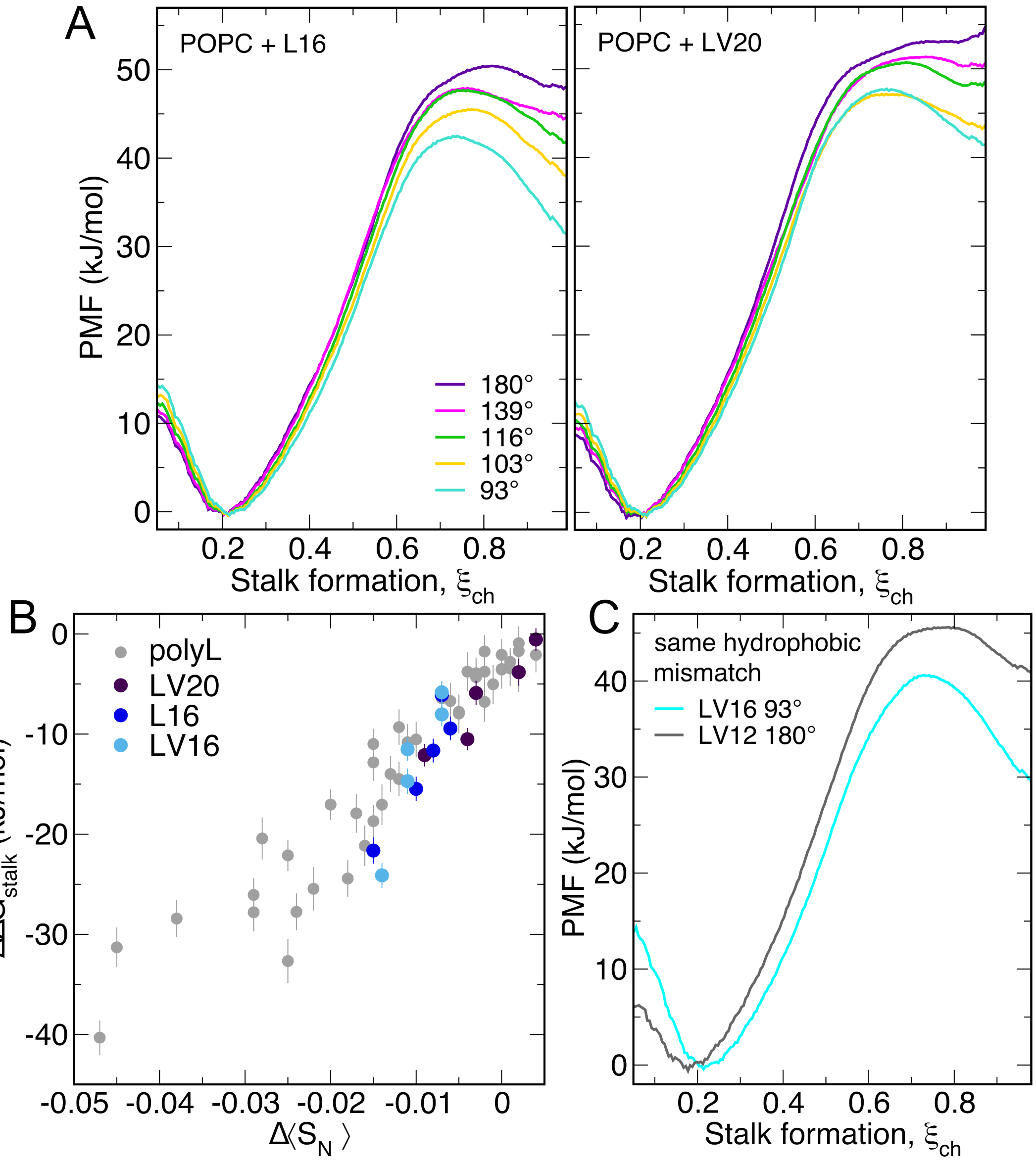


**Supplementary Figure S6:** (A) PMFs of stalk formation between two POPC bilayers with inserted L16 ($K_3W(LV)_{16}K_3$, left) or LV20 ($K_3W(LV)_{10}K_3$, right) with different bending angles. (B) Change in stalk free energy $\Delta\Delta G_{stalk}$ versus change in order parameter $\Delta\langle S_N\rangle$ upon insertion of one LV20 TMD ($K_3W(LV)_{10}K_3$, dark violet), one L16 TMD ($K_3WL_{16}K_3$, blue), or one LV16 TMD ($K_3W(LV)_8K_3$, light blue) per bilayer with varied bending angles. For reference, values obtained for polyleucine TMDs are shown as gray dots. (C) PMFs of stalk formation between two POPC membranes with kinked LV16 TMDs ($K_3W(LV)_8K_3$) or with straight LV12 TMD ($K_3W(LV)_6K_3$). The two TMDs yield the same hydrophobic mismatch, while the kinked LV16 TMD decreases the stalk free energy more compared to the straight LV12 TMD, demonstrating that kinks in TMDs favor stalk formation additionally to effects by negative hydrophobic mismatch.
